# Resistance profiling of *Aspergillus fumigatus* to olorofim indicates absence of intrinsic resistance and unveils the molecular mechanisms of acquired olorofim resistance

**DOI:** 10.1101/2021.11.16.468917

**Authors:** Jochem B. Buil, Jason D. Oliver, Derek Law, Tim Baltussen, Jan Zoll, Margriet W.J. Hokken, Marlou Tehupeiory-Kooreman, Willem J.G. Melchers, Mike Birch, Paul E. Verweij

## Abstract

Olorofim (F901318) is a new antifungal currently under clinical development that shows both in vitro and in vivo activity against a number of filamentous fungi including *Aspergillus fumigatus*. In this study we screened *A. fumigatus* isolates for intrinsic olorofim-resistant *A. fumigatus* and evaluated the ability of *A. fumigatus* to acquire an olorofim-resistant phenotype. No intrinsic resistance was found in 975 clinical *A. fumigatus* isolates. However, we found that isolates with increased olorofim MICs (> 8 mg/L) could be selected using a high number of conidia and olorofim exposure under laboratory conditions. Assessment of the frequency of acquired olorofim resistance development of *A. fumigatus* was shown to be higher than for voriconazole but lower than for itraconazole. Sequencing the *PyrE* gene of isogenic isolates with olorofim MICs of >8 mg/L identified various amino acid substitutions with a hotspot at locus G119. Olorofim was shown to have reduced affinity to mutated target protein dihydroorotate dehydrogenase (DHODH) and the effect of these mutations were proven by introducing the mutations directly in *A. fumigatus*. We then investigated whether G119 mutations were associated with a fitness cost in *A. fumigatus.* These experiments showed a small but significant reduction in growth rate for strains with a G119V substitution, while strains with a G119C substitution did not exhibit a reduction in growth rate. These in vitro findings were confirmed in an in vivo pathogenicity model.

**Importance:** Olorofim represents an important new treatment option for patients with difficult to treat invasive fungal infections, including triazole-resistant *A. fumigatus* infection. Our study provides insights into one resistance mechanism and the potential dynamics of olorofim resistance, which will help to prevent and manage resistance selection. Such insights are critical to antifungal stewardship and to safeguard its prolonged use in clinical practice.

## Introduction

It is estimated that around 250,000 people worldwide suffer from invasive aspergillosis annually (1). Patients at risk include those with neutropenia and in recent years cases have been increasingly observed in intensive care unit patients, including those with severe influenza and coronavirus infection (2–6). The triazoles voriconazole and isavuconazole are the recommended first line agents for the management of invasive aspergillosis (7, 8). The use of other registered antifungal agents such as liposomal amphotericin B is limited due to toxicity while the echinocandins exhibit lower efficacy against *Aspergillus* compared to the triazoles. Furthermore, the triazoles are currently the only agents that can be administered orally. However, the use of triazoles is threatened by the emergence of azole resistance in *Aspergillus fumigatus,* (9) which has now been reported globally (10, 11). Voriconazole resistance in *A. fumigatus* infection was associated with a near doubling of mortality at day 42 compared to voriconazole susceptible infection in patients that were treated with voriconazole (12). Azole resistance is mainly driven by environmental exposure of *A. fumigatus* to azole fungicides, which selects for fungicide resistance mutations that confer cross resistance to medical triazoles. In regions with environmental resistance, any patient may present with azole-resistant invasive aspergillosis that complicates diagnosis and successful therapy. Thus, there is an urgent need for new antifungal agents with efficacy against both azole-susceptible and azole-resistant *A. fumigatus* infection.

Olorofim (F901318) is a new antifungal currently under clinical development that shows activity against a number of filamentous fungi including *A. fumigatus*. It belongs to a new orotomide class of antifungals and acts by selective inhibition of fungal dihydroorotate dehydrogenase (DHODH), an essential enzyme within the *de novo* pyrimidine biosynthesis pathway (13, 14). The spectrum of activity of olorofim includes *Aspergillus* species including cryptic *Aspergillus* species and azole-resistant *Aspergillus* isolates, (15–22) *Lomentospora prolificans*, *Scedosporium* species (14, 23–27), agents of endemic mycoses such as *Coccidioides* species, *Histoplasma* species, *Sporothrix schenckii* and *Blastomyces* species (14, 28). Furthermore, in vitro studies show activity against *Madurella mycetomatis (29)*, *Microascus/Scopulariopsis*, *Penicillium*, *Paecilomyces*, *Purpureocillium*, *Rasamsonia* and *Talaromyces* species (14, 30). The *in vitro* activity was confirmed in *in vivo* models for pulmonary aspergillosis with azole-susceptible and azole-resistant isolates of *A. fumigatus* (14, 22), in a murine model of disseminated *A. terreus* aspergillosis (31), a murine model of chronic granulomatous disease infected with *A. fumigatus*, *A. nidulans*, and *A. tanneri (18)* and in a murine model of central nervous system coccidioidomycosis (28).

Antifungal drug resistance may develop through genetic variation that is created by the fungus to enable its adaptation to stress factors in its environment. Although drug exposure is not relevant to the development of resistance mutations per se, antifungal drug selection pressure is critical to create dominance of resistant cells within a population of fungal cells. The risk of resistance selection to a new class of antifungal drugs such as the orotomides will depend on the frequency of spontaneous mutations that confer an orotomide-resistant phenotype and the extent of selection pressure through patient therapy. In this study we screened for intrinsic olorofim-resistant *A. fumigatus*, evaluated the ability of *A. fumigatus* to develop an olorofim-resistant phenotype, and characterized underlying olorofim resistance mechanisms, including the effect of mutated DHODH on olorofim affinity and the impact of resistance mutations on virulence in a mouse model of disseminated aspergillosis. Lastly, the effect of mutated DHODH on conferring olorofim resistance in *A. fumigatus* was proven by introducing the mutation in a wildtype *A. fumigatus* strain.

## Results

### Agar based screening of resistance

A total of 976 clinical *A. fumigatus* isolates were screened for olorofim non-wildtypes on agar plates containing 0.125 mg/L olorofim. Only one isolate showed growth on the agar well supplemented with 0.125 mg/L olorofim (isolate V179-44). No growth on plates containing 0.125 mg/L olorofim was seen in the other 975 isolates.

### Identification of acquired resistance in A. fumigatus

Clinical *A. fumigatus* isolate (V179-44) was identified as possibly olorofim resistant and in vitro susceptibility testing using a spore suspension derived directly from the agar well supplemented with 0.125 mg/L of olorofim, showed an olorofim MIC of >8 mg/L. Susceptibility testing from the initial culture of strain V179-44 resulted in a wildtype olorofim MIC of 0.031 mg/L. Identification of the resistant isolate through beta-tubulin sequencing confirmed the conventional identification as *A. fumigatus* sensu strictu (32). As we suspected selection of a colony with a spontaneous olorofim resistance mutation, we tried to replicate this observation. Inoculation of >1×10^9^ spores of three *A. fumigatus* isolates (AZN8196, V052-35 and V139-36) on three 90 mm petri dishes with RPMI1620 agar supplemented with 2% glucose containing 0.5 mg/L olorofim, two olorofim-non-wildtype *A. fumigatus* colonies were retrieved from parental strain AZN8196 (AZN8196_OLR1 and AZN8196_OLR2), three from V052-35 (V052-35_OLR1 to V052_OLR3) and 11 from V139-36 (V139-36_OLR1 to V139-36-OLR11).

### In vitro frequency of spontaneous mutations resulting in olorofim resistance in asexual sporulation

Six *A. fumigatus* isolates (ATCC 204305, AZN8196, V052-35 (TR_34_/L98H), V139-36, V180-37 and V254-51 (Table 1) were used for the resistance frequency experiment. A total of 131 olorofim non-wildtype strains were retrieved, all of which showed an olorofim MIC of >8 mg/L (Table S1). An olorofim resistance frequency of 1.3 × 10^−7^ to 6.9 × 10^−9^ was observed (Figure 1). The mean itraconazole resistance frequency was between 1.2 × 10^−6^ and 3.3 × 10^−8^ and the mean voriconazole resistance frequency was between 1.8 × 10^−8^ and 2.0 × 10^−10^. Overall, the frequency of resistance was higher for itraconazole compared to olorofim, while voriconazole had the lowest frequency of resistance. The frequency of resistance of olorofim was significantly lower than itraconazole for strains AZN8196 and V254-51, while the frequency was not significantly lower for strains ATCC204305, V139-36 and V180-37. The frequency of voriconazole resistance was significantly lower compared with olorofim for strains ATCC204305, AZN8196, V139-36 and V254-51, while no significant differences were observed for strain V180-37. The second independent resistance frequency analysis using isolate Af293, which was cultured on yeast nitrogen base agar with glucose supplemented with 0.25 mg/L olorofim resulted in a mean frequency of olorofim resistance of 1.7× 10^−9^, a rate comparable to the above experiments. A total of 11 isolates were retrieved from parent strain Af293 of which 10 had an olorofim MIC of >8 mg/L while one had a MIC of 0.25 mg/L (Table S1).

**Table 1.** Strains used in this study for resistance frequency analysis.

| Strain | Cyp51A genotype | Olorofim MIC (mg/L) | Voriconazole MIC (mg/L) | Itraconazole MIC (mg/L) |
| --- | --- | --- | --- | --- |
| ATCC 204305 | wildtype | 0.016 | 0.25 | 0.125 |
| AZN8196 | wildtype | 0.031 | 0.25 | 0.125 |
| V052-35 | TR<sub>34</sub>/L98H | 0.031 | 8 | >16 |
| V139-36 | wildtype | 0.063 | 0.25 | 0.25 |
| V180-37 | wildtype | <0.016 | 0.5 | 0.5 |
| V254-51 | wildtype | 0.031 | 0.25 | 0.5 |
| AF293 | wildtype | 0.016 | 0.5 | 0.5 |

**Figure 1.**
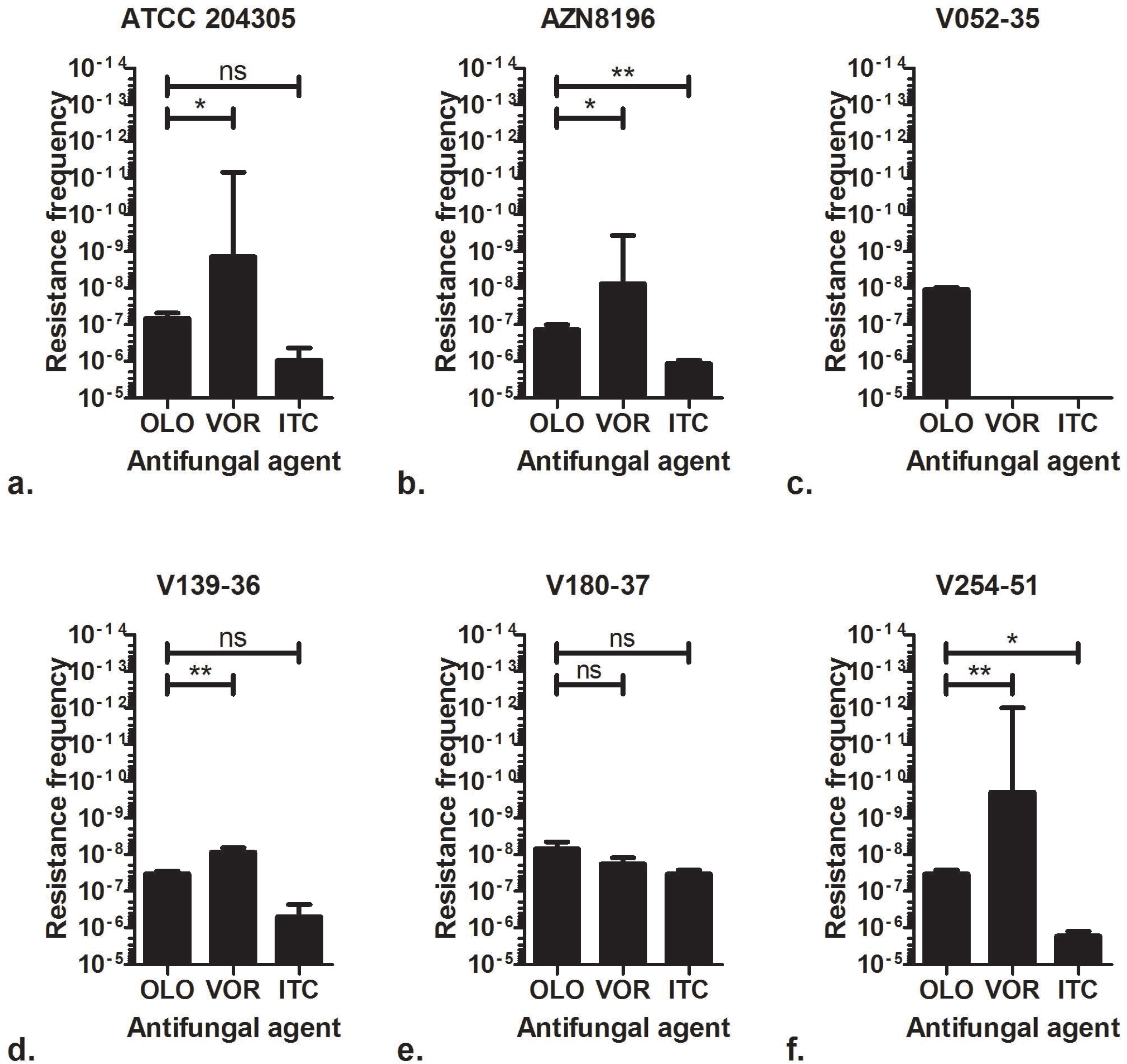
Olorofim resistance frequency. Frequency of resistance observed of six *A. fumigatus* isolates when 10^6^ to 10^9^ spores were incubated on RPMI agar plates containing either 0.5mg/L olorofim (OLO), 4 mg/L voriconazole (VOR) or 8 mg/L itraconazole (ITC). *A. fumigatus* ATCC 204305 b. *A. fumigatus* AZN 8196 c. *A. fumigatus* V052-35 (TR_34_/L98H, azole resistant) d. *A. fumigatus* V139-36 e. *A. fumigatus* V180-37 and f. *A. fumigatus* V254-51). * P=<0.05 ** P=<0.01, ns Not significant.

### Sequencing of pyrE identifies a hotspot for olorofim resistance at Gly119

The target of olorofim has been identified as the pyrimidine biosynthetic enzyme DHODH, which in *A. fumigatus* is encoded by the *pyrE* gene. When sequencing the full *pyrE* gene of olorofim strains retrieved from parent strain Af293, we found mutations at locus G119 in 10 of 11 sequenced olorofim-non-wildtype isolates. A single isolate with an olorofim MIC of 0.25 mg/L had a *pyrE* sequence identical to the parent Af293 strain. Subsequent analysis of a subset of 39 isolates from the resistance frequency analysis, showed that mutations that resulted in an amino acid substitution at G119 were present in 38/39 isolates. In total 7 isolates had a G119A amino acid substitution, while we found G119C (21 isolates), G119F (1 isolate), G119Y (1 isolates), G119S (11 isolates), and G119V (7 isolates) amino acid substitutions in the other isolates. One olorofim-resistant isolate harbored a H116P amino acid substitution in the PyrE gene (Table 2).

**Table 2.** Mutations in the *PyrE* gene in isolates selected for olorofim resistance.

| Parent Strain | Cyp51A genotype | Number of olorofim-resistant progeny sequenced strains | *PyrE* amino acid substitutions |  |  |  |  |  |  |
| --- | --- | --- | --- | --- | --- | --- | --- | --- | --- |
|  |  |  | G119A | G119C | G119F | G119Y | G119S | G119V | H116P |
| ATCC 204305 | wildtype | 8 | 1 | 5 |  |  | 2 |  |  |
| AZN8196 | wildtype | 6 | 2 | 2 |  |  | 1 | 1 |  |
| V052-35 | TR<sub>34</sub>/L98H | 4 |  | 4 |  |  |  |  |  |
| V139-36 | wildtype | 9 | 1 | 4 |  | 1 | 1 | 2 |  |
| V180-37 | wildtype | 4 | 1 | 1 | 1 |  |  |  | 1 |
| V254-51 | wildtype | 8 | 2 | 2 |  |  | 4 |  |  |
| Af293 | wildtype | 10 |  | 3 |  |  | 3 | 4 |  |

### Confirmation of the resistance mechanism

We investigated the effect that selected mutations at G119 had on the ability of olorofim to inhibit recombinant *A. fumigatus* DHODH. Recombinant DHODH with the amino acid substitutions G119A, G119V G119S and G119C showed significantly higher IC_50_ values for olorofim compared to wildtype DHODH (Figure 2). The substitutions at G119 thus result in decreased inhibition of *A. fumigatus* DHODH by olorofim, confirming the olorofim resistance mechanism.

**Figure 2.**
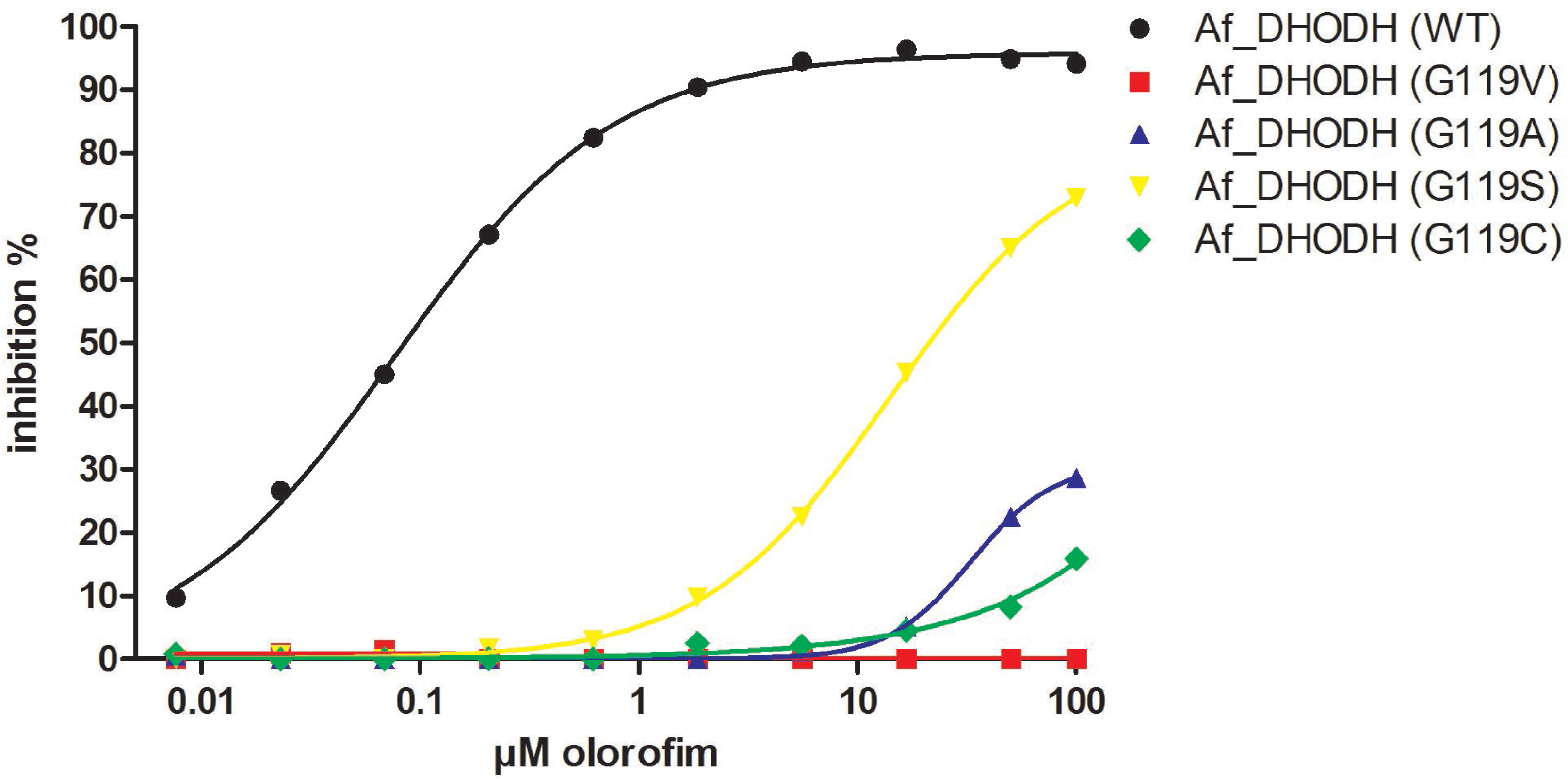
IC50s of wildtype and mutant DHODH. The inhibition of DHODH activity by a range of olorofim concentrations was measured for the recombinant wild type Af293 enzyme and the Gly119 mutants indicated. Lines were fitted using log(inhibitor) vs response – Variable slope (four parameters) in Graphpad Prism. R squares were 0.998 for Af_DHODH (WT), 0.556 for Af_DHODH (G119V), 0.924 for Af_DHODH (G119A), 1,000 for Af_DHODH (G119S) and 0.9680 for Af_DHODH (G119C).

### G119 transformations using CRISPR/Cas9

To further prove that the *PyrE* G119 mutations in *A. fumigatus* result in increased MICs to olorofim, we introduced the G119C mutation in *A. fumigatus* by a marker free CRISPR-Cas9 method in strain MFIG001 (33, 34). MFIG001 is a strain deficient in the non-homologous end-joining pathway resulting in a high transformation rate. Single colonies from the transformations were subcultured on Sabouraud dextrose agar (SDA) slants and screened for olorofim resistance on an agar plate containing 0.5 mg/L olorofim. Three strains (MFIG001_pyrE^g119c^_01, MFIG001_pyrE^g119c^_03 and MFIG001_pyrE^g119c^_05) that grew on the olorofim containing plate were selected for *PyrE* sequencing and subsequent MIC testing confirming the presence of the presence of G119C mutations (including the transformation specific synonymous PAM site mutation) (Figure S2) and olorofim MICs of >8 mg/L.

### Influence of PyrE substitution on radial growth rate

As development of resistance is often associated with attenuated virulence (35), we investigated the effects of olorofim resistance mutations on the fitness of *A. fumigatus*. To assess the impact of substitution of G119 in the *PyrE* gene on fitness, we used *in vitro* radial growth experiments. Mean growth curves are shown in Figure 3. The mean radial growth at day 5 of AZN8196_OLR1 (carrying the G119V mutation) was 30.7 mm, which was significantly different to the wildtype parent strain AZN8196 which had a mean growth of 40.0 mm (p<0.001). The mean radial growth of AZN8196_OLR2 (G119C) was 35.8 mm, also slightly reduced compared to strain AZN8196 but not significantly (P=0.06). The mean growth at day 5 was 42.7 mm for Af293, which was not significantly different compared with the mean growth at day of strain Af293_OLR7 (G119C), which had a mean growth 43.2 mm. The mean 5-day growth of Af293_OLR5 (G119S) was slightly decreased compared to parent at 39.5 mm (p=0.0039). Once more the glycine to valine mutation had the greater effect on growth with Af293_OLR9 growing 28.8 mm, 14.4 mm less that the parental strain (p=0.0010). Thus, in two different *A. fumigatus* strains the G119V mutants grew significantly more slowly than the parental strain (Figure 3).

**Figure 3.**
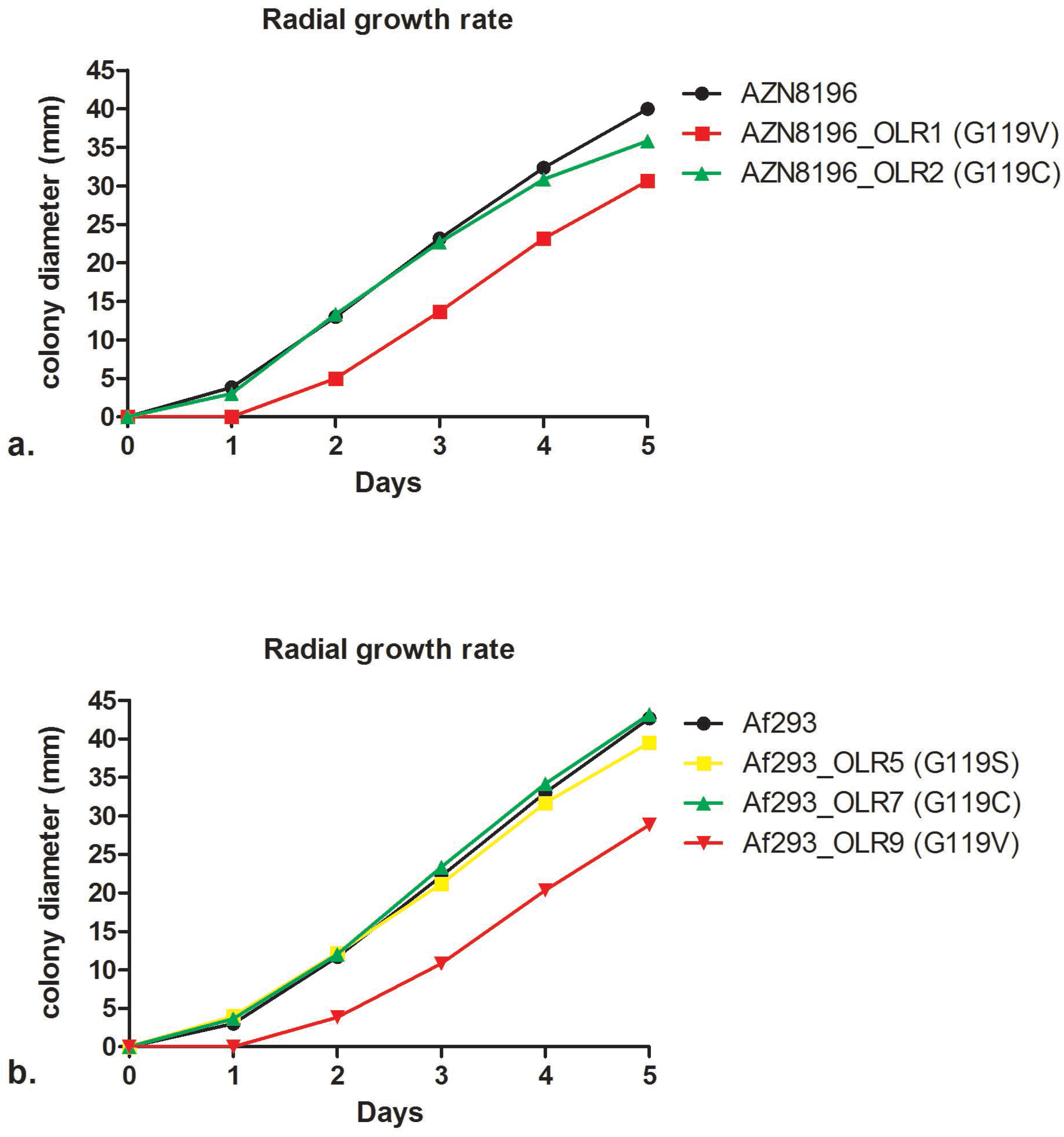
Radial growth rate of isolate AZN8196 and Af293 and olorofim resistant progeny. Colony diameters are displayed for a. isolate AZN8196 and 2 olorofim resistant progeny isolates AZN8196_OLR1 (G119V) and AZN8196_OLR2 (G119C) with and b. Af293, Af293_OLR5 (, Af293_OLR7 (G119S) Af293 OLR9 (G119V)

### Pathogenicity of olorofim-resistant A. fumigatus strains in an in vivo murine model

Although the radial growth rate experiments did not reveal significant fitness cost for two of the tested isolates, we wanted to confirm these observations in a neutropenic murine infection model. All strains demonstrated virulence with all animals succumbing to disease by 96 hours post infection. Median survival times of animals infected with AZN8196, AZN8196_OLR1 (G119V) and AZN8196_OLR2 (G119C) were 68.13, 89.75 and 73.38 hours after infection, respectively. Mice infected with AZN8196_OLR1 (G119V) survived significantly longer than those infected with their parent strain. There were no significant differences between mice infected with AZN8196_OLR2 (Figure 4a).

**Figure 4.**
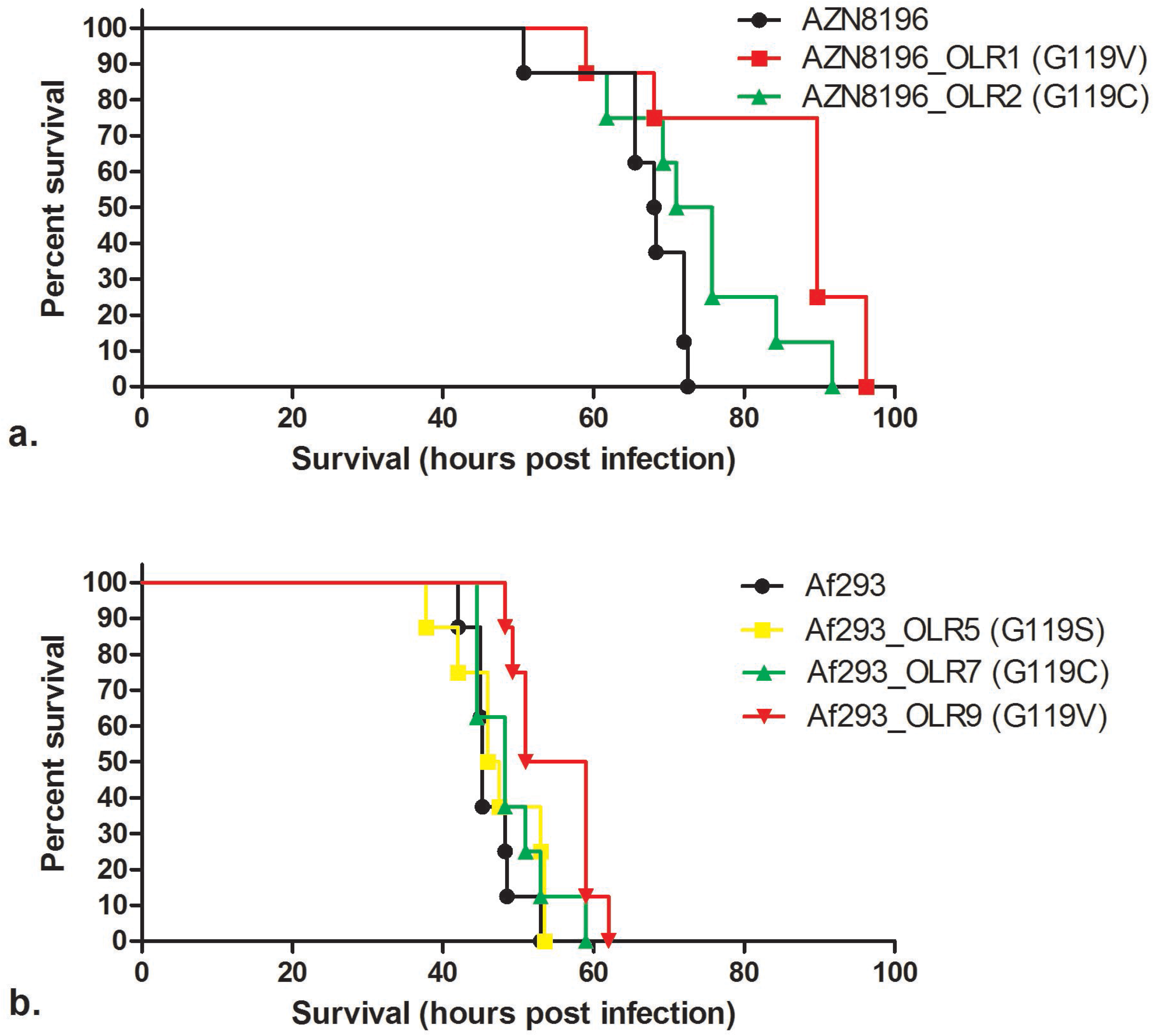
In vivo virulence model. Survival of mice inoculated with a. olorofim wildtype strain AZN8196 and olorofim resistant progeny AZN8196_OLR1 and AZN8196_OLR2, and b. olorofim wildtype strain Af293 and olorofim resistant progeny Af293_OLR5, Af293_OLR7 and Af293_OLR9. Eight mice were inoculated with each strain.

Median survival times of animals infected with Af293, Af293_OLR5 (G119S), Af293_OLR7 (G119C), and Af293_OLE9 (G119V) at a concentration of approximately 5 × 10^6^ CFU/mouse were 45.25, 46.75, 48.25 and 55 hours post infection, respectively. No difference in survival time was found between animals infected with Af293 and those infected with Af293_OLR5 (G119S) and Af293_OLR7 (G119C). However, Af293_OLR9 (G119V)-infected animals survived significantly longer than AF293-infected mice (Figure 4b).

Whilst the other mutants tested appeared as virulent as their parental strains, the two G119V mutant strains generated from different parents survived for longer. This is consistent with the slower growth observed for these strains in the radial growth experiments. These strains appear less fit than the wild type both *in vitro* and *in vivo*.

## Discussion

Evaluation of a large collection of clinical *A. fumigatus* isolates showed that the olorofim resistance frequency is negligible and no cross resistance with azoles was detected. However, olorofim resistance can be selected for under laboratory conditions and is associated with point mutations at locus G119 of the *PyrE* gene. Such mutations confer a resistant olorofim phenotype (olorofim MICs >8 mg/L), which appeared to have variable effects on virulence.

Olorofim is a promising novel antifungal with in vitro and in vivo efficacy against *A. fumigatus* infection, including triazole-resistant cases. The drug is currently undergoing phase II evaluation for treatment of patients with invasive fungal infections that cannot be managed with current agents. Screening of over 900 clinical *A. fumigatus* isolates showed intrinsic resistance is not identified, confirming the results from susceptibility testing of 1,032 clinical *A. fumigatus* isolates from Denmark (15).

Olorofim resistance in *A. fumigatus* has not been reported before. A previous evolution experiment involving 50 passages of *A. fumigatus* exposed to an olorofim concentration gradient, resulted only in a modest olorofim MIC increase. In contrast voriconazole generated a four-fold increase in MIC after only 15 passages (14). In the present study, we observed the *in vitro* acquisition of olorofim resistance while screening for intrinsic resistance using an agar supplemented with olorofim. All strains that were screened for olorofim resistance using this method were inhibited on this olorofim-containing agar, except one isolate. The olorofim-containing agar well of this strain showed growth of a single colony, in contrast to the growth control that showed confluent growth of numerous colonies on the whole agar surface. Had the initial isolate been resistant, we would have also expected numerous colonies growing in the olorofim-containing agar similar to the growth control. The lack of this growth, together with the discrepant results from the susceptibility testing of the parental colony and the colony growing on the olorofim-containing agar led us to believe that the resistant isolate had acquired olorofim resistance while being cultured on olorofim-containing agar.

By using a high inoculum of 1× 10^9^ CFU/mL we found that isolates with increased olorofim MICs (> 8 mg/L) could indeed be selected confirming our previous observation. As we wanted to understand the implications of this observation, we assessed the frequency of resistance development of *A. fumigatus* to olorofim and compared this frequency to other clinically used triazoles. We have chosen itraconazole and voriconazole as comparator agents as resistance development has been described in patients receiving long-term therapy for (cavitating) chronic pulmonary aspergillosis (CPA) but not in patients treated for acute invasive aspergillosis (36, 37). We found that the resistance frequency of itraconazole was higher than the resistance frequency found for voriconazole. Similar observations are seen in the treatment of patients with CPA where the rate of emergence of azole resistance during therapy was 13% for itraconazole and 5% for voriconazole (38). Differences in resistance frequency between itraconazole and voriconazole may be explained by the fact that almost all azole resistance associated substitutions reported in the *Cyp51A* gene result in itraconazole MICs above 4 mg/L, while only few substitutions result in high-level resistance to voriconazole (39). The finding of spontaneous olorofim resistance mutations is not surprising, but the frequency appears to be relatively low. The conditions that enable in vivo selection of olorofim resistance may be similar to those for triazole resistance; a setting of a high number of replicating fungal cells and chronic drug exposure. Such conditions may be present in patients with cavitary pulmonary lesions, such as aspergilloma and CPA but are unlikely in patients with acute invasive aspergillosis. However, as there are currently no alternative antifungal agents available for treatment of patients with triazole-resistant CPA that can be administered orally, olorofim represents a promising treatment option for this patient group that requires further clinical evaluation.

Importantly, *in vivo* selection of resistance mutations during treatment is not observed in patients treated for invasive aspergillosis. Triazole resistance in acute invasive aspergillosis is caused by inhalation of triazole-resistant *A. fumigatus* conidia that have developed resistance in the environment through exposure to azole fungicides, which occurred over several decades of exposure (40). Agents that inhibit DHODH as mode of action are currently not used for crop protection; to prevent a similar scenario to environmental triazole resistance selection, the use of similar mode of action compounds for medical and environmental applications should be avoided.

DHODH is an essential enzyme in the *de novo* pyrimidine biosynthesis pathway, and disruption of this pathway results in attenuated virulence in *A. fumigatus (41)*. Similar observations are reported in other fungal species like *Candida albicans, Cryptococcus neoformans* and *Histoplasma capsulatum* and the necessity of an undisrupted pyrimidine biosynthesis pathway is demonstrated in both *in vitro* and *in vivo* models (42–44). As DHODH, the product of the *pyrE* gene, is the enzyme target of olorofim action, we hypothesized that the most likely target of resistance is the *pyrE* gene (14). Indeed, sequencing the *PyrE* gene of isolates with olorofim MICs of >8 mg/L identified various amino acid substitutions. A homology model of the *A. fumigatus* DHODH predicted a potential binding mode for olorofim (14). Locus G119, located within this binding site was identified as a specific hotspot for olorofim resistance in *A. fumigatus* as 48/49 sequenced isolates had amino acid substitutions at G119. A single isolate had an amino acid substitution at position H116 which was predicted as a key residue for olorofim binding (14). The effect on olorofim susceptibility of mutations in *PyrE* at locus G119 was proven by both olorofim inhibition assay of mutant recombinant DHODH and by introducing the *PyrE* G119C mutation directly in *A. fumigatus*.

It remains uncertain whether locus G119 will also be the main mechanism of resistance if olorofim resistant *pyrE* isolates eventually emerge in clinical practice. However, similar in vitro resistance induction experiments were performed for triazole resistance in *A. fumigatus.* The mutations found in these in vitro experiments, like the amino acid substitutions at locus G54 and locus M220 in the *Cyp51A* gene can also be found in isolates retrieved from patients with CPA who are treated for long periods with triazoles, indicating that such in vitro experiments may predict the resistance mechanisms that can be found through clinical use (45, 46).

As development of antifungal resistance is often associated with attenuated virulence (35), we investigated whether amino acid substitutions in *pyrE* at locus G119 mutations were associated with a fitness cost in *A. fumigatus*. Analysis of *A. fumigatus* with disrupted *chsC* and *chsG* which encode Class III chitin synthases, showed a reduced colony radial growth rate compared to the wildtype strain. Subsequent assessment of pathogenicity in neutropenic mice showed a reduction in mortality in the mice inoculated with a *chsC* and *chsG* disrupted strain compared to the wildtype isolates (47). Similar correlations between growth rate and virulence were observed when the growth rate of *A. fumigatus* was assessed in 96-wells plates using the optical density as indicator for growth rate (48). To understand whether *PyrE* amino-acid substitutions influence fitness of *A. fumigatus* which may be extrapolated to *in vivo* pathogenicity, we analyzed the radial growth rate of five isolates with *PyrE* substitutions. These experiments showed a small but significant reduction in growth rate for strains with a G119V substitution (strain AZN8196_OLR1 and Af293_OLR9), while strains with a G119C substitution did not exhibit a reduction in growth rate. These in vitro findings were confirmed in the in vivo pathogenicity model whereas no significant difference in survival was observed for isolates with a G119C amino acid substitution (isolates AZN8196_OLR2 and Af293_OLR7). These results indicate that the amino acid substitution affects the binding of olorofim to DHODH but may not affect the function of DHODH itself and the effect on DHODH function is dependent on the underlying amino acid substitution. Furthermore, compensatory evolution has been shown to occur in triazole-resistant *A. fumigatus* isolates when cultured in azole-free conditions, indicating that a potential fitness cost can be overcome (49). However, population dynamics such as competition with other (wildtype) genotypes and selection pressure will ultimately determine which genotype will become dominant.

Olorofim represents an important new treatment option for patients with difficult to treat invasive fungal infections, including triazole-resistant *A. fumigatus* infection. Our study provides insights into one mechanisms and potential dynamics of olorofim resistance, which will help to prevent and manage resistance selection in various patient groups. Such insights are critical to antifungal stewardship and to safeguard its prolonged use in clinical practice.

## Material and Methods

### Agar based screening of resistance

We screened 976 clinical *A. fumigatus* isolates that were cultured between 2015 and 2017 for non-wildtype olorofim phenotypes. Inoculum with a density of approximately 0.5 McFarland was prepared in sterile 0.9% NaCl with 0.1% Tween 20 and one drop of 25 μl was used to inoculate an agar plate (RPMI1640 with 2% glucose) containing 0.125 mg/L olorofim. An agar plate containing only RPMI1640 with 2% glucose agar was used as growth control. Olorofim MIC-testing was performed on isolates growing on the agar plate containing olorofim. If routine susceptibility results indicated resistance to voriconazole or itraconazole, the *Cyp51A* gene was subsequently sequenced.

### Minimal inhibitory concentration of olorofim

Susceptibility testing was performed using the EUCAST method for susceptibility testing of molds E.Def.9.3.1 (EUCAST.org). Olorofim pure powder was obtained from F2G (Manchester, United Kingdom). Stock solutions of olorofim were prepared in DMSO. 96-wells plates with 2-fold dilutions of olorofim were prepared in RPMI1680 with 2% glucose and buffered with MOPS. The olorofim concentration range used was 0.016–8 mg/liter. Inoculum was prepared in sterile 0.9% NaCl with 0.1% Tween 20. Spores were harvested from mature culture and the suspension was adjusted to 80-82 % transmission at 530 nm (Spectrofotometer Genesys 20) to create a 1 - 4.2 × 10^6^ CFU/ml spore suspension (50). Inocula were added to the 96-well plates to create a final concentration of 2 – 5 × 10^5^ CFU/ml in each well. The inoculated plates were incubated for 48 h at 35 °C. MIC was defined as the lowest concentration without visible growth.

### Selection of resistant mutants

Six *A. fumigatus* isolates (ATCC 204305, AZN8196, V052-35 (TR_34_/L98H), V139-36, V180-37 and V254-51) were used for the olorofim resistance induction experiment. Sabouraud dextrose broth containing chloramphenicol (SAB-c) was inoculated and cultures were grown at 28°C. Spores were harvested in sterile saline with 0.05% tween 20 and the inoculum was transferred to a sterile vial. The spore suspension was adjusted to 1×10^9^ spores/mL using an hemocytometer. One mL spore suspension was added to a 90 mm agar plate containing RPMI 1640 +2% glucose (1.5% agar) containing 0.5 mg/L olorofim. Cultures were grown at 30°C. Isolates that grew on the olorofim containing plates were subcultured on SAB-c for subsequent MIC testing and DNA isolation

### Frequency of resistance analysis

Six *A. fumigatus* isolates (ATCC 204305, AZN8196, V052-35 (TR34/L98H), V139-36, V180-37 and V254-51 were used for the olorofim resistance induction experiment (Table 2). SAB-c was inoculated using a single spore isolated from the six parent strains and cultures were grown at 28°C. Spores were harvested in sterile saline with 0.05% tween 20 and the inoculum was transported to a sterile vial. The spore suspension was adjusted to 1×10^9^ spores/mL using an hemocytometer. One mL spore suspension was added to a 90 mm agar plate containing RPMI 1640 +2% glucose (15% agar) containing either 0.5 mg/L olorofim, 4 mg/L voriconazole or itraconazole 8 mg/L. These concentrations were chosen as these were the concentrations which are 2 dilutions higher that the concentration that inhibits 100% of wildtype *A. fumigatus* isolates (51). Cultures were grown at 30°C. Isolates that grew on the olorofim containing plates were subcultured on SAB-c for subsequent MIC testing and DNA isolation. The resistance rate was calculated by dividing the number of retrieved resistant colonies by the number of inoculated spores and the mean of 5 experiments was used for comparison. Differences in resistance frequency between olorofim and itraconazole or voriconazole were tested for significance using the student T test. Statistical significance was defined as a *P* value of ≤0.05 (two-tailed). To confirm the resistant rates, a second experiment was performed in another laboratory. Spore stocks of *A. fumigatus* strain Af293 were prepared and inoculated onto yeast nitrogen base with glucose agar (YNBG) containing 0.25 mg/L olorofim. A total of 8 × 10^9^ spores were inoculated into 12 × 100 ml YNBG-OLO agar plates that were subsequently incubated for 5 days at 35°C. Colonies growing on drug-containing plates were subcultured on YNBG-OLO to confirm resistance.

### Sequencing of PyrE identifies hotspot at Gly119

The *PyrE* gene of all isolates from parent strain Af293 were sequenced as previously described using primers AFDseq-F2 and AFDseq-R2 (15). *PyrE* amino acid sequences of olorofim-resistant strains were compared to the amino acid sequence of the wildtype parent strains. As these and the earlier pilot experiments showed only amino acid substitutions at locus *G119* without mutations at other loci in *PyrE*, we sequenced only part of the *PyrE* gene for the other strains. This part of the *A. fumigatus PyrE* gene was sequenced using primer PyrE_G119_Fwd: AGTAAAGGAGGCACCCAAGAAAGCTGG and PyrE_G119_Rev: GCCAATGGGGTTGTTGAGCGTATACCC. We randomly selected 39 olorofim-resistant strains from the resistant frequency analysis.

### Olorofim inhibition assays of mutant recombinant DHODH

The cloning of *A. fumigatus* DHODH_(89-531)_ cDNA into protein expression vector pET44 yielding pET44AFD was described previously (14). For preparation of mutated protein this plasmid was mutated at codon 119 using the Phusion Site-Directed Mutagenesis kit (Thermo Scientific). PCR reactions were set up with Phusion HSII polymerase, pET44AFD as a template, with one constant primer (AFDSDM_R1; CCTCTTCCGCGTCGGGATAA) and a variable that had a single codon change (CGCATCATATT*xyz*GTGGAAGCTCT). The sequence of *xyz* (GGT in wild type) was: GTT for G119V; GCT for G119A; AGT for G119S; TGT for G119C. The PCR product representing a linear version of pET44AFD with the mutation present was ligated using T4 DNA ligase and transformed into Max Efficiency DH5α competent cells (Thermo Fisher). Sequencing confirmed the desired mutations were present. The constructs were transformed into *E. coli* BL21 (DE3) cells (Merck) and the mutant proteins were expressed and purified according to the protocol described by Oliver, et al (14). DHODH assays were set up in the presence and absence of olorofim at concentrations between 0.008 to 100 μM. Assays were carried out in 50 mM Tris HCl pH8, 150 mM KCl, 10% (wt/vol) glycerol, and 0.1% (wt/vol) Triton X-100 in the presence of 1 mM L-dihydroorotic acid, 0.05 mM coenzyme Q2 and 0.1 mM 2,6-dichloroindophenol as a redox indicator. The reaction was followed by absorbance at 600 nm and reaction velocities used to construct IC50 curves. (14). Curves were fitted in GraphPad Prism using variable slope (four parameters) on log transformed data.

### G119 transformations using CRISPR-Cas9

To prove that G119 mutations are resulting in increased olorofim MICs we introduced the G119C mutation in strain MFIG001 (34) as previously described (33). In short, protoplasts were generated by inoculation of approximately 1 × 10^6^ fresh conidia in Yeast extract Glucose medium (YG; 0.5% yeast extract, 2% glucose) for 16 hours at 37 °C shaking at 120 rpm. Mycelia were harvested by filtration and resuspended in YG with protoplasting buffer (5 g Vinotaste in 50 mL 1M KCl and 0.1 M Citric Acid added to 50 mL YG) and reincubated for 4 hours at 37 °C shaking at 100 rpm. Then protoplast were washed 3 times in 0.6 M KCl and 50 mM CaCl_2_ and the protoplast concentration was adjusted and diluted to approximately 1 × 10^6^ protoplasts. A PAM site close to the G119 locus was selected with an adjacent 20 nucleotide protospacer that covers the G119 locus using a web-based guide RNA designing tool EuPaGDT (doi:10.1099/mgen.0.000033). The genome sequence of *A. fumigatus* A1163 (Aspergillus_fumigatusa1163.ASM15014v1) was manually uploaded to EuPaGDT to design gRNAs to the *pyrE* locus. The program was carried out with the default settings and the crRNA with the highest QC score closest to the target integration was selected for transformation. A single stranded (ss) DNA repair template was selected that covered 50 nucleotides on both sites of the protospacer and PAM site (Figure S1) that contains a synonymous point mutation in the PAM site and the T>G mutation in the G119 locus resulting in GGT (Glycine) to TGT (Cysteine) transformation. Alt-R^®^ CRISPR-Cas9 tracrRNA, Alt-R^®^ CRISPR-Cas9 crRNA, Alt-R^®^ S.p. Cas9 Nuclease and ssDNA repair template were ordered from Integrated DNA Technologies. Ribonucleoprotein (RNP) complex were assembled *in vitro* in Nuclease-Free IDTE buffer. RNP complex, the ssDNA repair template and PEG buffer ((60% wt/vol PEG3350, 50 mM CaCl2, 450 mM Tris-HCl, pH 7.5) were mixed with 50 μl of the 1 × 10^6^ /ml protoplast and incubated on ice for 50 minutes. Then 1 mL of PEG 3350 was added to the solution and incubated for another 25 minutes at 20 °C. The solution was incubated on five yeast extract peptone dextrose (YPD) agar plates and incubated for 48 hours. Single colonies were subcultured on SDA slants and screened for resistance by spotting 2 μL containing 100-500 conidia on RPMI1680 2% glucose agar containing 0.125 mg/L olorofim and RPMI1680 2% glucose agar without olorofim as growth control. Five isolates showing prominent growth after 48 hours were selected and further analyzed by olorofim MIC testing and sequencing of the *pyrE* G119 hotspot as described earlier.

### Radial growth rate

To study the radial growth rate we randomly selected five olorofim-resistant isolates, two from strain AZN8196 (AZN8196_OLR1, AZN8196_OLR2) and three from strain Af293 (Af293_OLR5, Af293_OLR7, and Af293_OLR9) from the initial induction experiments and compared the radial growth rate to the wildtype parent strain. The strains were chosen as these strains harbored the most common amino acid substitutions G119C, G119V and G119S. We assessed the growth by measuring the colony diameters in the horizontal axis and vertical axis once every 24 hours for a period of 5 days. We used an initial inoculum of 2 × 10^2^ spores, quantified with an hemocytometer for all strains and inoculated the spores in the middle of a 90mm petri dish with Yeast Nitrogen Base with glucose. We did three independent experiments per strain and reported the mean of the three experiments. Differences in growth rate in mm/day at day 5 between wildtype and olorofim-resistant strains were tested for significance using the student T test. Statistical significance was defined as a *P* value of ≤0.05 (two-tailed).

### Assessment of the pathogenicity of olorofim-resistant progeny compared to strain AZN8196 and Af293

To study the virulence of isolates with *PyrE* amino-acid substitution, we assessed survival of these isolates in a murine model of disseminated aspergillosis and compared the survival to their wildtype parent strains. Ideally, truly isogenic isolates are used for such experiments. However, the selected isolates were selected for olorofim resistance on a plate and only the *PyrE* gene was sequenced and we thus cannot exclude amino acid substitutions elsewhere in the genome. To exclude effects of such additional substitutions we used five separately selected olorofim-resistant strains to perform the experiments. CD-1 mice (Charles River Laboratories, Margate, UK) were immunosuppressed 3 days prior to infection with cyclophosphamide administered at 200mg/kg subcutaneously. Inoculum was prepared for *A. fumigatus* strains AZN8196, AZN8196_OLR1, AZN8196_OLR2, Af293, Af293_OLR5, Af293_OLR7, and Af293_OLR9. Mice were infected by intravenous administration of 0.2mL conidial suspension. An inoculum of approximately 5 × 10^5^ CFU/mL was used for strains AZN8196, AZN8196_OLR1, AZN8196_OLR2 and an inoculum of approximately 2 5 × 10^7^ CFU/mL was used for strains Af293, Af293_OLR5, Af293_OLR7, and Af293_OLR9 resulting in 1 × 10^5^ CFU/mouse and 5 × 10^6^ CFU/mouse respectively. These inocula were chosen as those are the LD_90_ doses that were previously determined for these specific *A. fumigatus* strains (48, 52). The concentration of conidia was adjusted using a hemocytometer and confirmed by quantitative culture on SDA. Actual and intended inoculum levels are listed in table S1. Eight mice were inoculated with either strain. Mice were monitored for survival for 10 days and euthanized when they demonstrated high weight loss, signs of sepsis or severe torticollis. Survival data was analyzed using GraphPad Prism (Version 5.3) and checked for significance using the Log-rank (Mantel-Cox) Test. Statistical significance was defined as a *P* value of ≤0.05 (two-tailed).

---

**Table S1.**
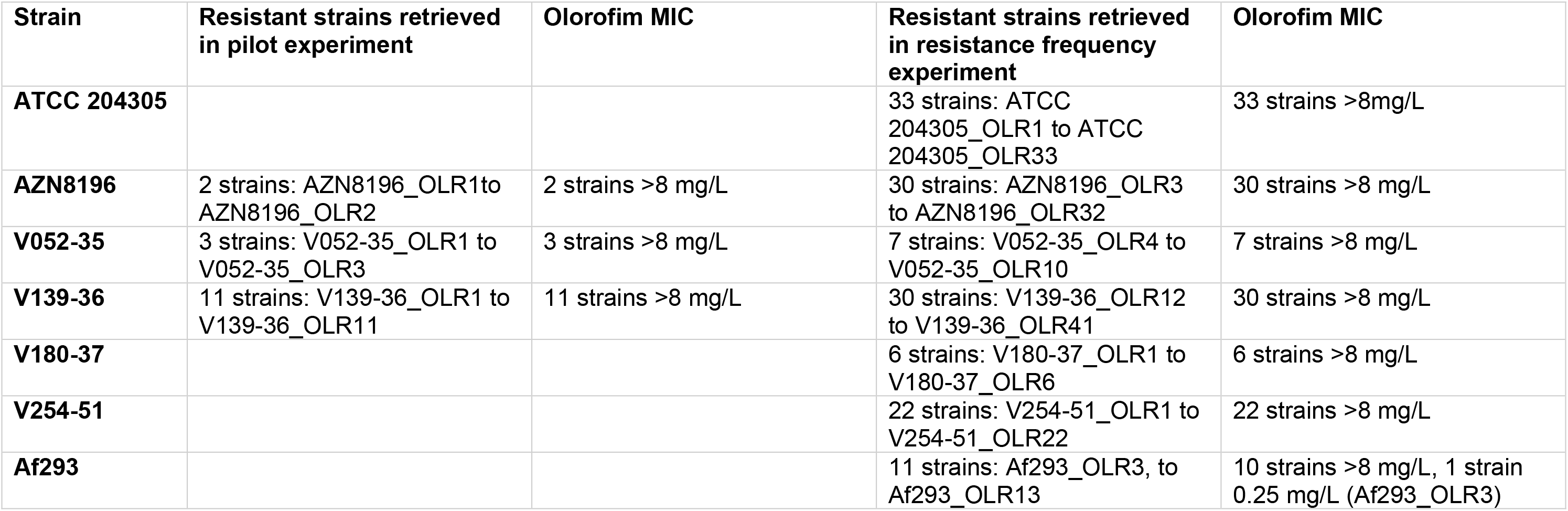
Olorofim resistance progeny strains and corresponding MIC.

| Strain | Resistant strains retrieved in pilot experiment | Olorofim MIC | Resistant strains retrieved in resistance frequency experiment | Olorofim MIC |
| --- | --- | --- | --- | --- |
| **ATCC 204305** |  |  | 33 strains: ATCC 204305_OLR1 to ATCC 204305_OLR33 | 33 strains >8mg/L |
| **AZN8196** | 2 strains: AZN8196_OLR1to AZN8196_OLR2 | 2 strains >8 mg/L | 30 strains: AZN8196_OLR3 to AZN8196_OLR32 | 30 strains >8 mg/L |
| **V052-35** | 3 strains: V052-35_OLR1 to V052-35_OLR3 | 3 strains >8 mg/L | 7 strains: V052-35_OLR4 to V052-35_OLR10 | 7 strains >8 mg/L |
| **V139-36** | 11 strains: V139-36_OLR1 to V139-36_OLR11 | 11 strains >8 mg/L | 30 strains: V139-36_OLR12 to V139-36_OLR41 | 30 strains >8 mg/L |
| **V180-37** |  |  | 6 strains: V180-37_OLR1 to V180-37_OLR6 | 6 strains >8 mg/L |
| **V254-51** |  |  | 22 strains: V254-51_OLR1 to V254-51_OLR22 | 22 strains >8 mg/L |
| **Af293** |  |  | 11 strains: Af293_OLR3, to Af293_OLR13 | 10 strains >8 mg/L, 1 strain 0.25 mg/L (Af293_OLR3) |

**Table S2.** *Aspergillus fumigatus* strains used in the *in vivo* virulence model

| Strain | PyrE amino acid substitution | Intended inoculum (CFU/mL) | Intended inoculum (CFU/mouse) | Actual inoculum (CFU/mL) | Actual inoculum (CFU/mouse) | Median survival time (hours post infection) | Statistical comparison to <sup>1</sup>AZN8196 or <sup>2</sup>Af293 |
| --- | --- | --- | --- | --- | --- | --- | --- |
| **AZN8196** | - | 5 x 10<sup>5</sup> | 1 x 10<sup>5</sup> | 5.05 x 10<sup>5</sup> | 1.01 x 10<sup>5</sup> | 68.13 | - |
| **AZN8196_OLR1** | G119V | 5 x 10<sup>5</sup> | 1 x 10<sup>5</sup> | 4.10 x 10<sup>5</sup> | 8.20 x 10<sup>4</sup> | 89.75 | P=0.0065<sup>1</sup> |
| **AZN8196_OLR2** | G119C | 5 x 10<sup>5</sup> | 1 x 10<sup>5</sup> | 3.40 x 10<sup>5</sup> | 6.80 x 10<sup>4</sup> | 73.38 | NS<sup>1</sup> |
| **Af293** | - | 2.5 x 10<sup>7</sup> | 5 x 10<sup>6</sup> | 2.15 x 10<sup>7</sup> | 4.30 x 10<sup>6</sup> | 45.25 | - |
| **Af293_OLR5** | G119S | 2.5 x 10<sup>7</sup> | 5 x 10<sup>6</sup> | 2.05 x 10<sup>7</sup> | 4.10 x 10<sup>6</sup> | 46.75 | NS<sup>2</sup> |
| **Af293_OLR7** | G119C | 2.5 x 10<sup>7</sup> | 5 x 10<sup>6</sup> | 2.55 x 10<sup>7</sup> | 5.10 x 10<sup>6</sup> | 48.25 | NS<sup>2</sup> |
| **Af293_OLR9** | G119V | 2.5 x 10<sup>7</sup> | 5 x 10<sup>6</sup> | 1.35 x 10<sup>7</sup> | 2.70 x 10<sup>6</sup> | 55.00 | P=0.0022<sup>2</sup> |

CFU: colony forming units. NS: not significant

**Figure S1.**
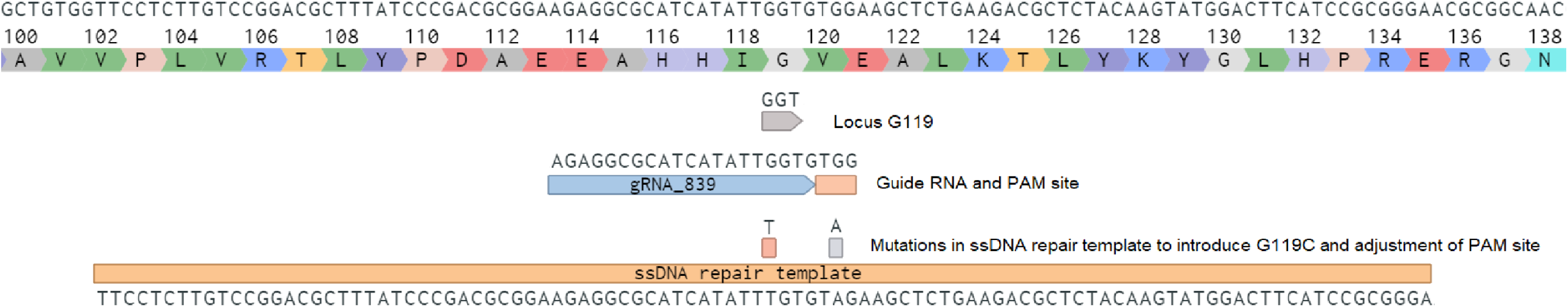
Details of CRISPR-Cas9 components for introducing the G119C mutation in A. fumigatus PyrE gene. Amino acid 100 to 138 of the PyrE gene are shown. The hotspot region G119, the guide RNA (AGAGGCGCATCATATTGGTG) and PAM site (TTG) are annotated. Furthermore the mutations in the single stranded DNA repair template compared to WT *PyrE* sequence are noted. The G>T in locus G119 results in formation of Cysteine. The G>A mutation in the PAM site leads to a synonymous mutation. This mutation results in the incapability of binding of the ribonucleoprotein complex due to disruption of the PAM site.

**Figure S2.**
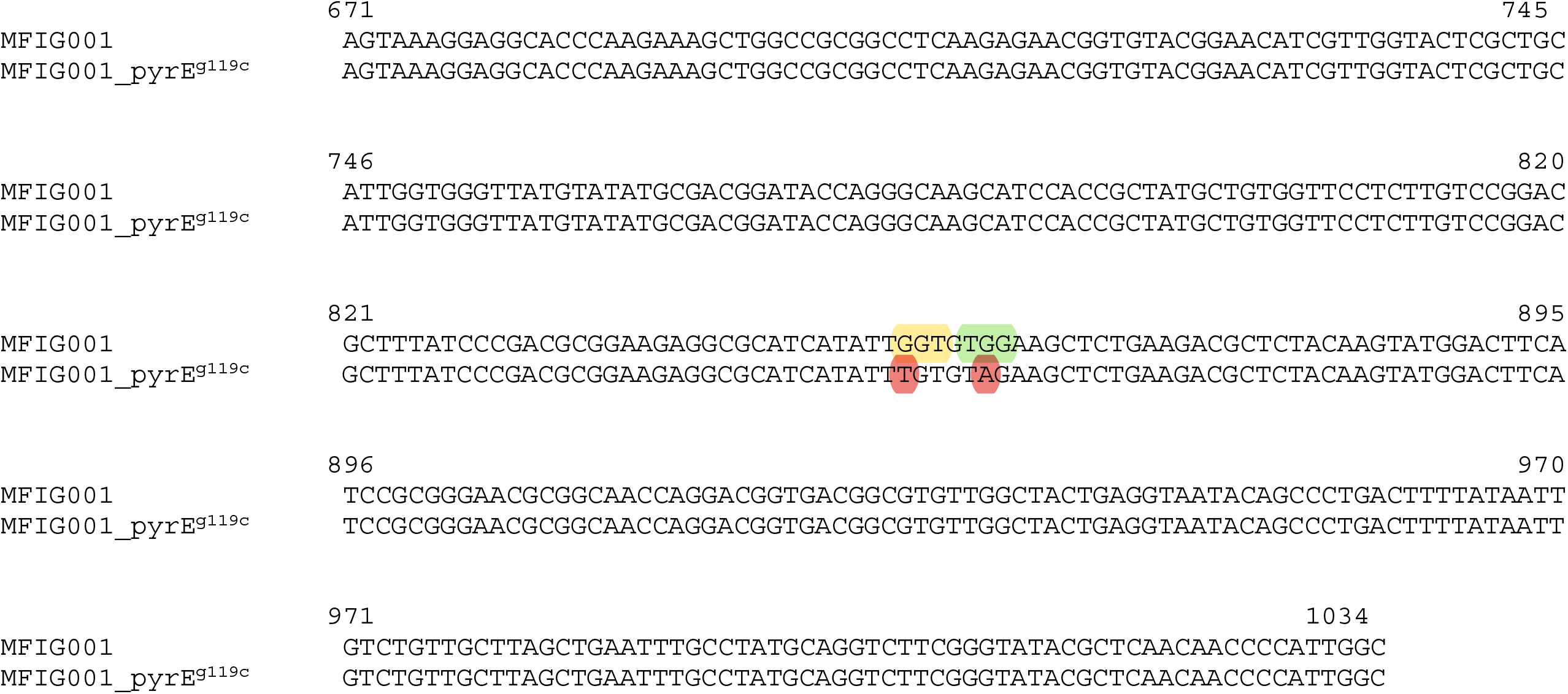
Alignment of *PyrE* sequence of the *pyrE* wildtype MFIG001 strain and the MFIG001_pyrE^g119c^_01, MFIG001_pyrE^g119c^_03 and MFIG001_pyrE^g119c^_05 strains. The G119 locus is marked in yellow. The PAM site is marked in green and the introduced G119C mutation and synonymous PAM site mutations are marked in red.

